# Neck-region–microtubule interactions direct counterclockwise stepping of kinesin-1

**DOI:** 10.1101/2025.06.07.658466

**Authors:** Song-Ho Chong, Ryota Iino

## Abstract

The “neck” region of kinesin is a structurally conserved element critical for force generation, stepping directionality, and cargo transport along microtubules, yet its atomic-scale structure in a functional context remains unresolved. Here, we employ all-atom replica exchange molecular dynamics simulations to resolve a high-confidence conformation of the neck region in dimeric human kinesin-1 bound to a realistic microtubule lattice, and use this structure to simulate kinesin’s initial stepping motion. Our simulations reveal that the neck coiled-coil is oriented perpendicular to the microtubule’s axis and positioned near its surface—a conformation consistent with earlier proposals but lacking high-resolution validation. Importantly, simulations indicate that neck–microtubule interactions bias the stepping trajectory, directing the rear kinesin head to overtake the front head from the right (counterclockwise stepping). These findings establish a mechanism by which neck–microtubule interactions govern directional bias in kinesin’s initial step, offering new insight into the molecular basis of its motility.

## INTRODUCTION

Kinesins are molecular motor proteins that transport intracellular cargo along microtubules, playing critical roles in various cellular processes such as mitosis, organelle trafficking, and neuronal transport.^1–3^ Among them, kinesin-1 (hereafter referred to as kinesin), which is the primary focus of the present study, represents a prototypical member of the kinesin family known for its high processivity and well-characterized plus-end-directed motility. These motors convert chemical energy derived from ATP hydrolysis into mechanical work, enabling highly processive, stepwise movements along microtubule tracks. Kinesin typically functions as a dimer, and its motility is driven by the coordinated actions of its two motor heads, which alternately bind to and release from the microtubule in a hand-over-hand fashion. ^4,5^ The co-ordination and unidirectionality of this stepping motion depend critically on conformational changes within specific structural elements of kinesin, notably the “neck” region.

The neck region of kinesin comprises two structurally well-conserved segments located at the C-terminal end of the motor head: the flexible neck linker of ∼12 amino acids and the subsequent neck helix of ∼30 amino acids. The neck linker undergoes conformational transitions, switching from an undocked, disordered state to a docked, *β*-strand conformation upon ATP binding to the head.^6–9^ This conformational change plays a pivotal role in force generation and forward stepping of the kinesin motor. In contrast, the neck helix—a short *α*-helical segment following the neck linker—is essential for dimerization of the two motor heads via formation of a coiled-coil structure,^10^ which in turn plays an important role in governing the processive run length.^11^ Although this coiled-coil motif has been confirmed in solution-phase X-ray crystal structures,^10^ its precise conformation and possible interactions with microtubule in the functionally relevant, microtubule-bound state remain uncharacterized at atomic resolution. This lack of high-resolution structural information hampers a comprehensive understanding of how the neck region contributes to kinesin stepping, particularly through its potential interactions with the microtubule surface.

Here, we investigate the structural and functional roles of the human kinesin neck region using all-atom molecular dynamics (MD) simulations. Due to the lack of high-resolution experimental structures for the neck region connecting the two heads (see Figure 1), we first determine its conformation in the presence of the microtubule using the generalized replica exchange with solute tempering (gREST) method, ^12^ a variant of replica exchange MD implemented in the GENESIS software.^13,14^ gREST enhances conformational sampling by selectively applying temperature scaling to a user-defined solute region—here, the neck-linker region—while maintaining the rest of the system at physiological temperature.^15–18^ This allows for efficient modeling of elusive segments within large biomolecular assemblies such as the kinesin–microtubule complex, without disrupting their global structural integrity. To ensure the robustness of our findings and reduce force-field dependence, we perform gREST simulations using two widely adopted force fields—AMBER ff99SB-ILDN ^19,20^ and CHARMM36m^21^—and compare the results. In both cases, the neck coiled-coil consistently adopts a conformation perpendicular to the microtubule’s long axis and closely associated with its surface. Using this structurally refined model, we then simulate the initial stepping motion of kinesin to investigate how interactions between the neck region and the microtubule influence directional dynamics. Our results reveal that these interactions strongly bias the motion of the rear kinesin, overtaking the front kinesin from the right side (counterclockwise stepping). Together, these findings provide mechanistic insight into how neck-region– microtubule interactions direct the trajectory of kinesin’s initial stepping motion, offering a new structural perspective on kinesin motility.

**Figure 1:**
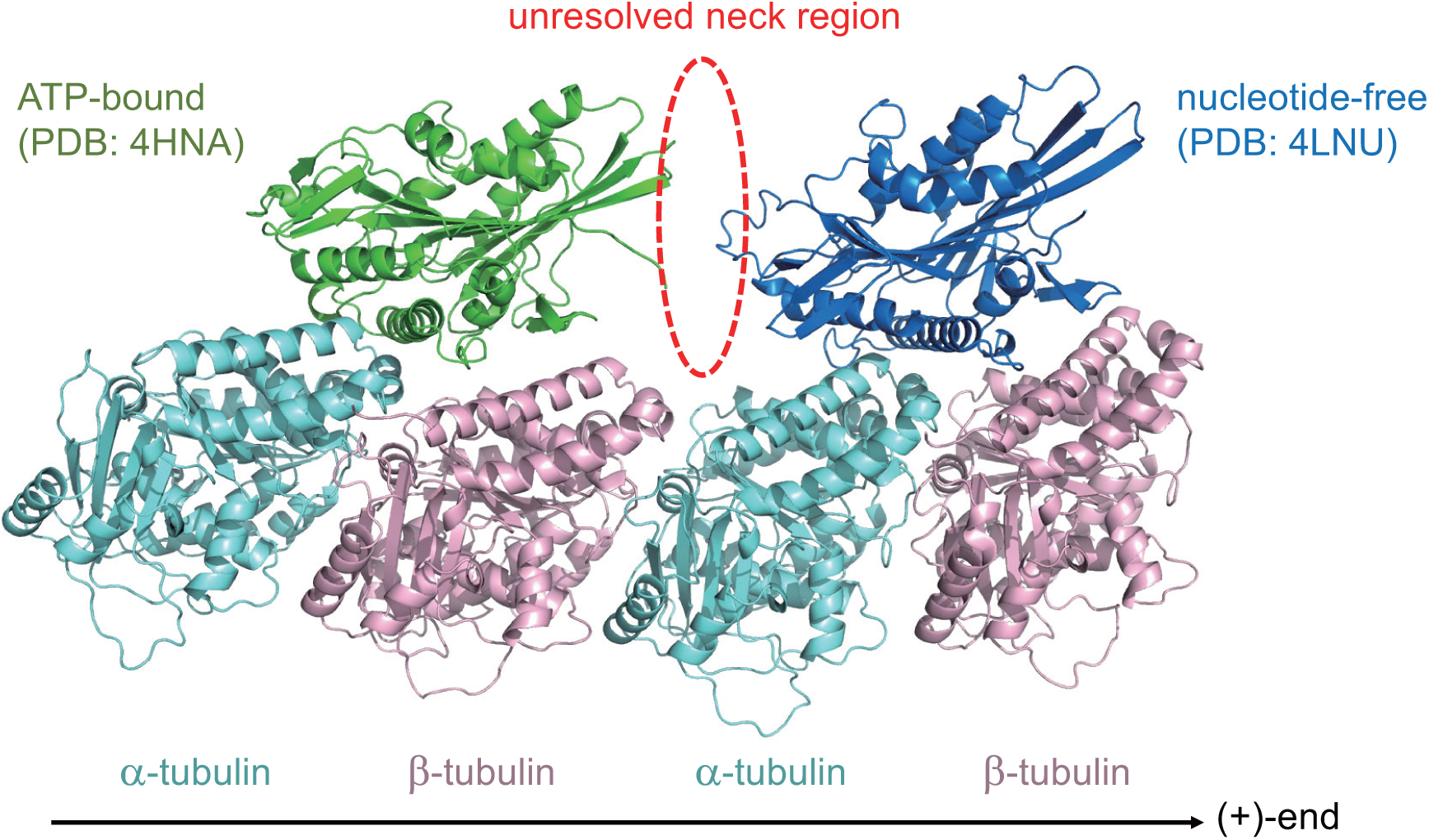
Kinesin dimer on a microtubule based on available experimental structures. The PDB entry 4HNA^23^ was used for the ATP-bound rear head positioned on the *αβ*-tubulin heterodimer subunit, while the PDB entry 4LNU^24^ was employed for the nucleotide-free front head on the *αβ*-tubulin subunit. These tubulin subunits were connected using the microtubule structure (PDB entry 3J6F ^25^) as a template. The unresolved neck region connecting the two heads is indicated by a dashed red oval.

## RESULTS

### Initial modeling of the neck region

To perform MD simulations of the dimeric kinesin on a microtubule, we first constructed an initial structural model, with particular attention to the neck region. Given the absence of a complete high-resolution structure of human kinesin-1 in the microtubule-bound state, we adopted a hybrid modeling strategy: wherever experimentally determined atomic-resolution structures were available, we incorporated them directly; for missing segments, we used homology or *ab initio* modeling via MODELLER.^22^

For the ATP-bound rear kinesin head, we used the X-ray structure of PDB entry 4HNA,^23^ which includes a docked conformation of the neck linker. For the nucleotide-free front kinesin head, we employed PDB entry 4LNU.^24^ Both X-ray structures also contain a single *αβ*-tubulin heterodimer subunit to which the kinesin head is bound, and this subunit was used to guide the proper placement of both kinesin heads on the full microtubule lattice (Step 1 in Figure 2). To model the microtubule, we adopted the high-resolution structure of PDB entry 3J6F,^25^ which comprises three protofilament rows, each consisting of three repeating *αβ*-tubulin heterodimers. This extended, realistic multi-protofilament arrangement was chosen to provide a structurally accurate landing site for kinesin stepping simulations and to capture potential lateral interactions between the kinesin neck region and the microtubule surface— features not represented in simplified one-dimensional models. By contrast, the so-called E-hooks—C-terminal tails of *α*- and *β*-tubulins that are disordered and rich in negatively charged residues^26^—were not included in our model. These regions are absent from available PDB structures due to their intrinsic flexibility, and modeling them alongside the already challenging neck region would have substantially increased the system’s complexity. We will discuss the potential effects associated with the E-hooks in the Discussion section.

**Figure 2:**
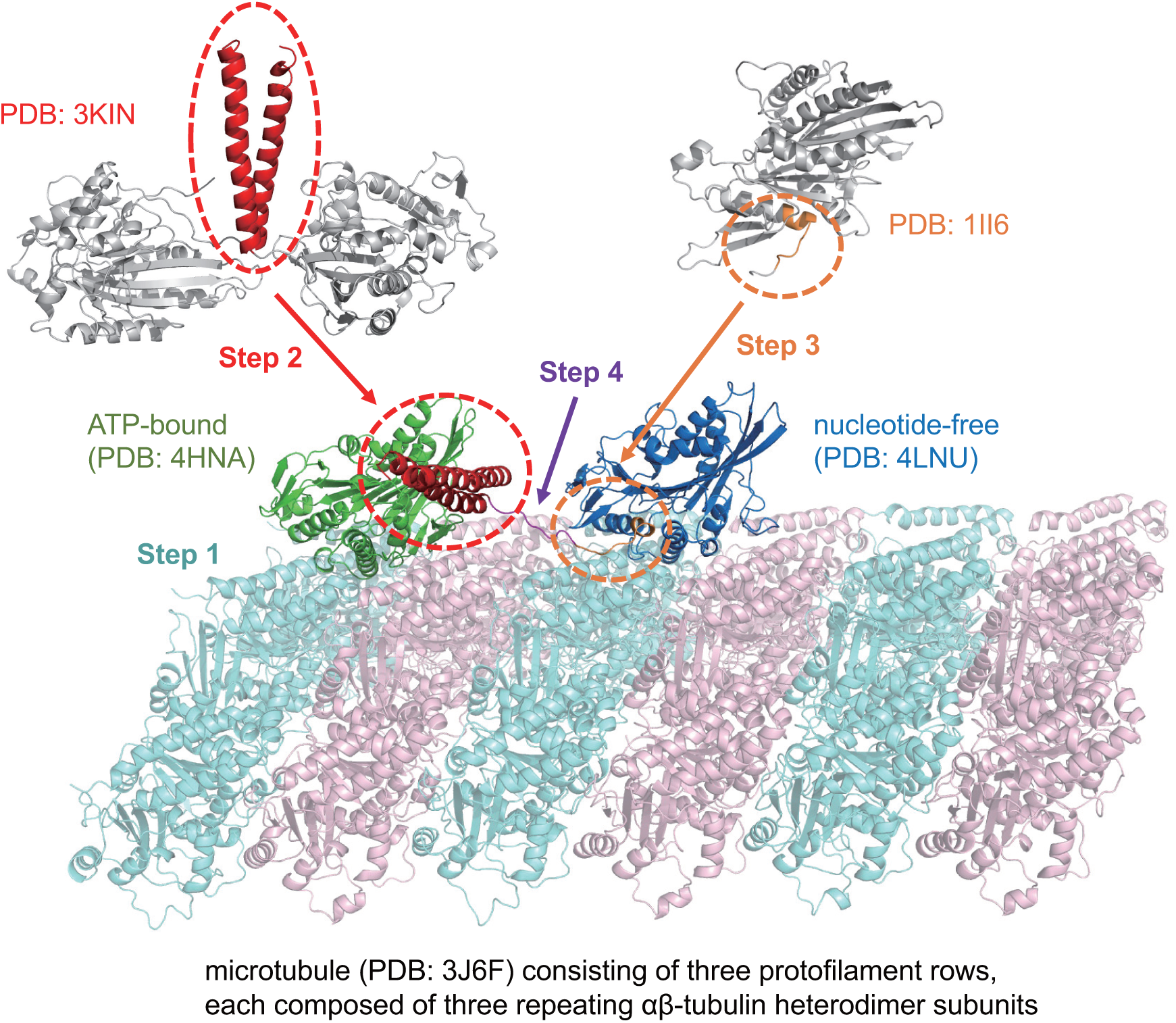
Initial modeling of the neck region of dimeric kinesin on a microtubule. Step 1: ATP-bound rear head (PDB: 4HNA^23^) and nucleotide-free front head (PDB: 4LNU^24^) were placed onto a microtubule model consisting of three parallel protofilaments (PDB: 3J6F^25^). Step 2: The neck helices forming a coiled-coil structure were taken from the solution-phase dimeric structure of rat kinesin (PDB: 3KIN^10^) and used to model the neck helices of both kinesin monomers. Step 3: The undocked, rearward-extending neck linker from Eg5 kinesin (PDB: 1II6^27^) was used to model the front kinesin’s neck linker. Step 4: The remaining residues connecting the neck linker to the neck helix in the front kinesin were modeled using MODELLER. Further details of these steps are provided in the main text. The final model is shown in side and top views in Figure S1.

Modeling of the neck region began with the rear kinesin from PDB entry 4HNA,^23^ which contains the head and a docked neck linker but lacks the downstream neck helix. To complete this segment, we incorporated the solution-phase dimeric structure of rat kinesin (PDB entry 3KIN^10^), which contains both the docked neck linker (residues 325–336 in human kinesin-1) and the subsequent neck helix (residues 337–368). By aligning the overlapping neck linker segments from both structures, we generated a continuous model that seamlessly connects the motor head, neck linker, and neck helix (Step 2 in Figure 2). In this 3KIN dimer, the neck helix forms a coiled-coil with its counterpart from the second kinesin subunit. We therefore adopted the neck helix from the second chain to model the front kinesin’s neck helix, yielding a preformed coiled-coil arrangement that defines the spatial constraints of the dimeric neck conformation (Step 2 in Figure 2).

To complete the front kinesin’s neck region, we next modeled the neck linker segment connecting the motor head to the previously placed neck helix. In contrast to the rear kinesin, whose neck linker adopts a docked conformation extending toward the microtubule plus end, the neck linker of the front kinesin is undocked and extends rearward (toward the minus end) to engage in the coiled-coil interface. To construct this rearward-directed conformation, we used the undocked neck linker from Eg5 kinesin (PDB entry 1II6^27,28^) as a structural template (Step 3 in Figure 2). As this fragment did not fully span the distance to the neck helix, the remaining residues were modeled using MODELLER in *ab initio* mode (Step 4 in Figure 2). Together, these steps produced a complete structural model of dimeric human kinesin-1 (residues 1–370 for each monomer) bound to a microtubule (the final model is shown in side and top views in Figure S1), serving as the starting conformation for subsequent modeling with gREST.

### gREST simulations for the neck region

The accuracy of the initial structure just constructed remains limited, particularly in the neck region where high-resolution experimental data are lacking. To refine the conformation of the neck region, we performed enhanced conformational sampling using gREST.^12^ By exchanging temperatures only within the solute region—here, the neck-linker region—while keeping the rest of the system at physiological temperature, gREST enables efficient exploration of conformational space in the region of interest without disrupting the overall structural context. Since structural sampling of flexible regions such as the neck linker can be sensitive to the choice of force field, we performed gREST simulations using two widely used force fields: AMBER ff99SB-ILDN^19,20^ and CHARMM36m.^21^ In the following, we will mainly refer to results based on AMBER ff99SB-ILDN unless stated otherwise, and results based on CHARMM36m are reported in the Supporting Information.

To reduce the computational cost associated with gREST simulations, we constructed a smaller system (see Figure S2A) compared to the one used for the initial modeling shown in Figure 2. In the initial kinesin–microtubule model, the microtubule consisted of three protofilament rows, each containing three *αβ*-tubulin heterodimer subunits arranged longitudinally along the protofilament axis. This extended structure was intended for use in our future “walking” simulations, in which the rear kinesin steps over the front kinesin and requires a landing site. However, the distal *αβ*-tubulin subunits are unnecessary when focusing solely on conformational sampling of the neck region. Therefore, to reduce system size and computational load, we removed the final *αβ*-tubulin subunits from the microtubule (Figure S2A) and carried out gREST simulations on this truncated model.

Conformational sampling of the neck region during the gREST simulation was monitored by tracking the tip positions of the neck helix, which are indicated as red dots in Figure S2B. We performed 1 *µ*s gREST simulations and visualized the distribution of the red dots over time to assess sampling coverage. As shown in Figure S2B, we compared the spatial distributions of the red dots using data from the entire simulation (0–1000 ns) and from the later portion (200–1000 ns), excluding the initial 200 ns. The comparison revealed that structures sampled during the first 200 ns were primarily influenced by the initial model and were not revisited during the remaining 800 ns of simulation. This suggests that the initial 200 ns contain artifacts originating from the starting structure, and thus, we excluded this period from subsequent analyses and used only the data from 200 ns onward.

To identify representative conformations of the neck region, we performed clustering analysis based on the gREST simulation data. For the three-dimensional coordinates of the neck helix tip sampled during the 200–1000 ns interval, principal component analysis (PCA) was first applied to reduce the dimensionality of the positional data to two principal components. Subsequently, *k*-means clustering was performed in the PCA-reduced space to classify the sampled conformations. The results are shown in Figure 3A. We found that the conformational ensemble can be broadly divided into two distinct clusters, to be referred to as Cluster 1 and Cluster 2. Figure 3A also shows the density distribution of the sampled conformations, revealing that Cluster 1 is more frequently sampled than Cluster 2. Closer inspection of the density map reveals that Cluster 2 can be further subdivided into approximately five sub-clusters. However, these sub-clusters adopt similar conformations, and we therefore did not pursue detailed analysis of them. Representative structures corresponding to the center of each cluster are illustrated in Figure 3B, providing insight into the characteristic conformations adopted by the neck region.

**Figure 3:**
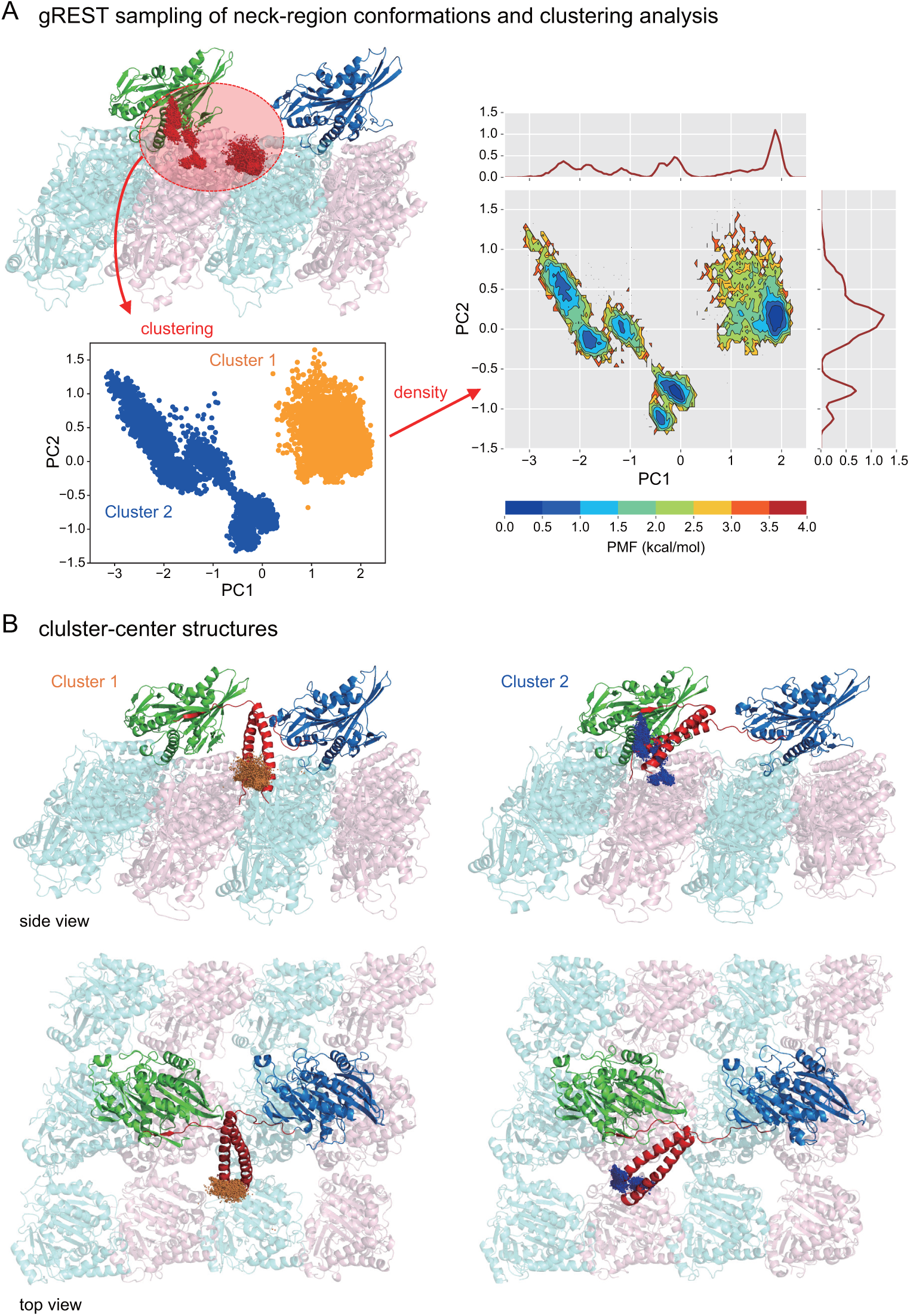
gREST sampling of neck-region conformations and clustering analysis. (A) Red dots indicate the tip positions of the neck helix sampled during the gREST simulations (see also Figure S2). These positions were projected onto the first two principal component (PC) axes, and *k*-means clustering was performed in this reduced space. Two major clusters were identified and labeled as Cluster 1 and Cluster 2. The right panel shows the potential of mean force (PMF) surface constructed from the sampling density in this reduced space. (B) Representative structures of Cluster 1 (left) and Cluster 2 (right) in side and top views.

The results described thus far were obtained using the AMBER force field (ff99SB-ILDN^19,20^). To assess force field dependence, we performed the same analysis on the gREST simulation data obtained with the CHARMM force field (CHARMM36m^21^). The results are summarized in Figure S3. We found that, similar to the AMBER case, the conformational ensemble of the neck region can be broadly divided into two major clusters. The most densely populated cluster obtained with CHARMM (Cluster 1’ in Figure S3B) closely corresponds to Cluster 1 from the AMBER results. In this conformation, the neck helix is oriented perpendicular to the microtubule’s long axis and positioned near the microtubule surface. The consistent emergence of the Cluster 1 conformation as the dominant structural state across both force fields underscores its reliability as a representative model of the neck region.

We note, however, that in the 1 *µ*s gREST simulation performed with the CHARMM force field, the microtubule structure became partially destabilized. Specifically, the distance between the central protofilament and one of its neighboring protofilaments increased significantly, leading to partial disruption of the overall microtubule architecture (Figure S4, right). This structural instability was not observed in the AMBER simulations (Figure S4, left). For this reason, in the following walking simulations, we used only the AMBER force field to ensure structural integrity of the microtubule throughout the simulation.

### Initial stepping motion of the kinesin dimer on microtubule

To investigate how the identified neck-region conformations influence kinesin motion, we performed MD simulations starting from the representative cluster structures. Given the substantial computational cost required to simulate the full kinesin walking cycle, we focused on the initial stage of the stepping motion, in which the rear kinesin detaches from the microtubule and advances approximately half a step forward. Simulations of the complete walking cycle are currently in progress.

Simulations of kinesin stepping require the presence of a landing site on the microtubule. We therefore restored the distal *αβ*-tubulin subunits removed during the gREST simulations and used this extended microtubule as the starting structure (Figure S5A and B). To promote detachment of the rear kinesin head, we replaced ATP with ADP, which is known to reduce kinesin’s binding affinity to the microtubule. ^29^ We then carried out 20 independent simulations, each lasting several hundred nanoseconds. However, spontaneous detachment was not observed in any trajectory, suggesting that much longer timescales may be required beyond what is feasible with our available computational resources. Recently, nucleotide-dependent conformational changes of the kinesin head bound to the microtubule have been reported,^30^ and such conformational changes may be required for detachment from the microtubule.

In light of this limitation, we adopted a more drastic yet conceptually motivated approach to promote detachment of the rear kinesin. Specifically, we removed the *αβ*-tubulin subunit located directly beneath the rear kinesin head, along with neighboring subunits from adjacent protofilaments, creating a local void beneath the head (Figure S5C). Detachment from the microtubule is thought to be governed by two main factors: the nucleotide state of the head^29,31^ and the internal strain between the two heads.^30,32,33^ Because our model uses ADP-bound kinesin, which has inherently weak microtubule affinity, and removes the underlying tubulin, the setup allows us to specifically examine how internal strain transmitted through the neck linker contributes to the initiation of stepping. This provides a simplified yet controlled framework for probing the mechanical role of the neck region in kinesin’s initial stepping.

Using the modified setup, we conducted 20 independent simulations, each starting from the Cluster 1 conformation in Figure 3B. In many trajectories, the rear head advanced toward the microtubule plus end within approximately 100 ns. Representative cases are shown in Figure 4 and Figure S6. To aid visualization, the removed tubulin subunits were reinserted in the figures to illustrate the full structural context. Each figure displays structural snapshots at 0 ns and 120 ns (Figure 4A and B; Figure S6A and B), while the center-of-mass positions of the rear head sampled every 1 ns during 1–120 ns are plotted as green spheres to illustrate its intermediate movement (Figure 4C; Figure S6C). In Figure 4, the rear head moves through the right side of the front head; in Figure S6, it moves over the top. To characterize overall trends in stepping direction, we compiled the center-of-mass positions of the rear head (green spheres; sampled every 1 ns) from all 20 simulations into a composite plot (Figure 5A). Despite variability across individual trajectories, a clear pattern emerged: when viewed from above, the rear head consistently followed a right-side, counterclockwise path relative to the front head. This directional bias suggests that conformation of the neck region and interhead tension via the neck region influence the early trajectory of stepping.

**Figure 4:**
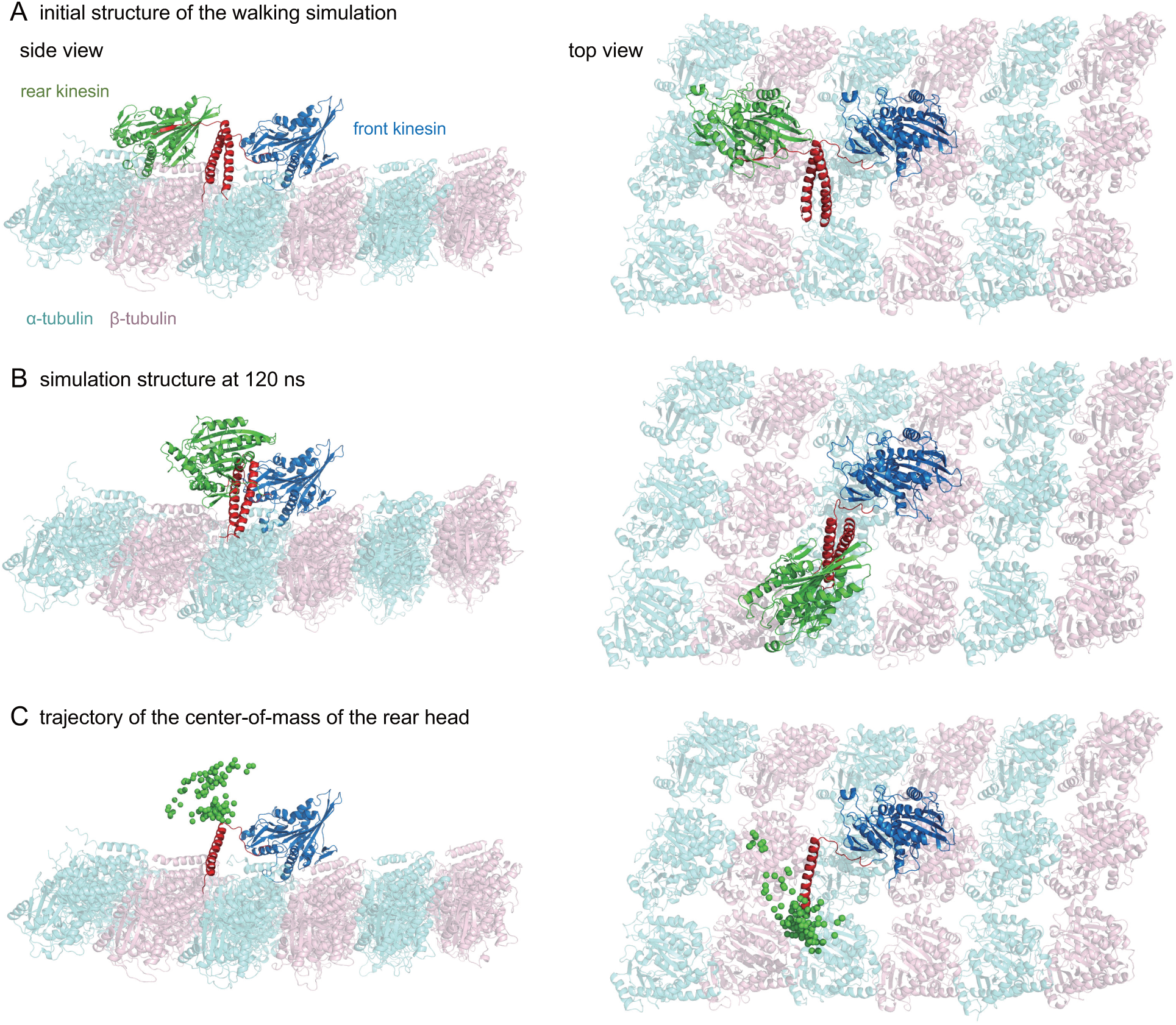
Representative trajectory of the initial stepping motion of the kinesin dimer on the microtubule. (A, B) Structural snapshots of the system at 0 ns and 120 ns, respectively, from a simulation initiated from the Cluster 1 conformation (see Figure 3B). To illustrate the full structural context, the *αβ*-tubulin subunits that were removed in the simulation setup have been reinserted. (C) Intermediate movement of the rear head is visualized by plotting its center-of-mass positions as green spheres, sampled every 1 ns from 1 to 120 ns.

**Figure 5:**
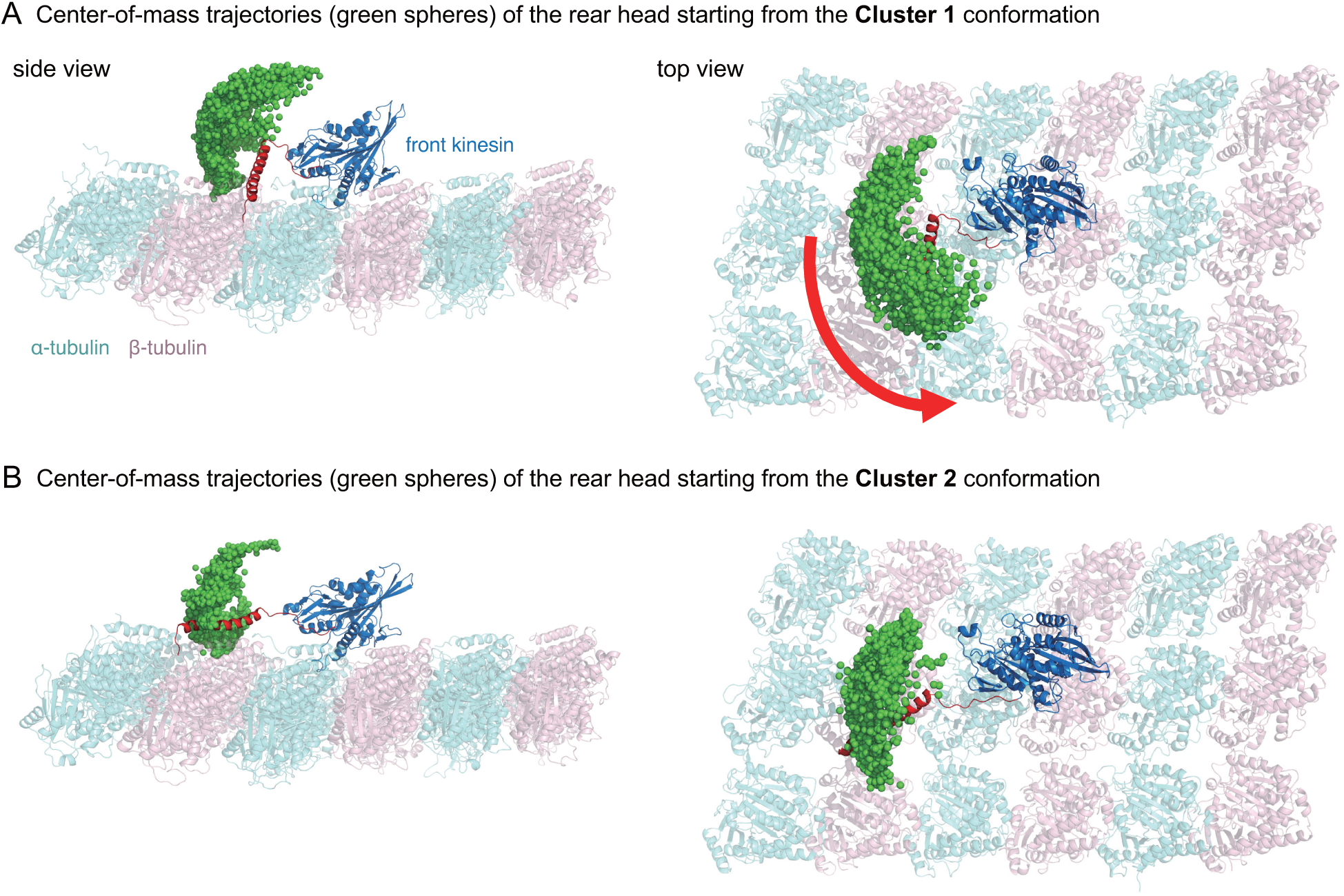
Composite analysis of kinesin stepping trajectories from simulations initiated with two distinct neck-region conformations. (A) Center-of-mass positions of the rear head (green spheres), sampled every 1 ns over the 120 ns duration of each of the 20 simulations starting from the Cluster 1 conformation (see Figure 3B). (B) Corresponding plot from the 20 simulations initiated with the Cluster 2 conformation.

To test the effect of an alternative neck conformation, we performed 20 simulations using the Cluster 2 structure in Figure 3B as the starting point. In this conformation, the neck region folds back and only weakly interacts with the microtubule surface. Despite removing the underlying tubulin subunits, the rear head did not move forward in any trajectory (Figure 5B). This contrast with Cluster 1 underscores that strain accumulation in the neck region alone is insufficient for productive stepping; it also requires interactions between the neck region and the microtubule. These findings highlight the critical role of neck-region–microtubule interactions in establishing the directional bias of kinesin’s initial stepping motion.

## DISCUSSION

This study presents an unprecedented all-atom MD investigation of the dimeric human kinesin-1 bound to the microtubule. While previous all-atom MD studies have examined monomeric kinesin bound to a single *αβ*-tubulin heterodimer subunit,^34–41^ to our knowledge, no prior work has reported simulations of a kinesin dimer interacting with a multi-protofilament segment of the microtubule. In particular, the neck region that is critical for directional stepping has never before been structurally modeled and simulated on the microtubule lattice in its functional context. Our system includes a realistic segment of the microtubule composed of three parallel protofilaments, providing a structurally accurate landing site and lateral interactions that are absent in simplified one-dimensional microtubule models. This allowed us to observe not only the initial stepping motion of the rear kinesin but also the directionality bias induced by interactions between the neck region and the microtubule surface. The resulting simulation system comprises approximately three million atoms when solvated, representing one of the largest MD simulations of a cytoskeletal motor protein reported to date.

The neck region of kinesin is a structurally unresolved but functionally essential element. By applying gREST simulations, we determined a reliable conformation of the neck region on a realistic microtubule lattice. The gREST method, ^12^ which enables enhanced conformational sampling by selectively tempering only the particular region of interest, proved particularly effective for exploring the flexible and structurally elusive neck region. The most representative structure obtained from our gREST simulations (Cluster 1 conformation in Figure 3B) shows that the neck coiled-coil is oriented perpendicular to the microtubule’s long axis and positioned close to the microtubule surface. Importantly, this conformation was consistently observed regardless of the force field used in the gREST simulations—both AMBER ff99SB-ILDN^19,20^ and CHARMM36m^21^ yielded essentially the same result—supporting the robustness of the structure. Strikingly, this predicted conformation closely resembles an early structural model proposed in Ref. 42 (see Figure 4 in Ref. 42) and a recent subnanometer-resolution 3D reconstruction of kinesin dimer on microtubule^43^ (see Figure 3 of Ref. 43). This agreement between atomistic simulations and experimental models reinforces the plausibility of the identified neck-region conformation.

We found that such a neck-region conformation that allows interactions with the microtubule surface plays a critical role in determining the directionality of kinesin’s initial stepping motion. Indeed, this conformation consistently guides the rear kinesin to move along counterclockwise trajectories relative to the front kinesin during the early phase of stepping (Figure 5A). Notably, this predicted counterclockwise motion aligns well with prior experimental observations. High-speed/high-precision dark-field microscopy reported that, upon detachment from the microtubule, the rear kinesin tends to fluctuate at the right-hand side of the front kinesin,^44,45^ consistent with counterclockwise movement. A similar lateral deviation was also observed using high-speed/high-precision interferometric scattering microscopy.^46^ More recently, optical trapping experiments revealed a transient intermediate state in which the rear kinesin aligns laterally beside the front kinesin before completing the forward step,^47^ although in the structural model proposed in Ref. 47, the neck region is oriented perpendicular to the microtubule surface and interactions between these two components are not assumed. Additionally, interferometric MINFLUX microscopy showed that, upon entering the one-head-bound intermediate state, the rear head localizes to the right side of the front head.^48^ Our simulations thus not only provide atomistic support for these experimental findings, but also offer mechanistic insight into how neck-region–microtubule interactions introduce directional bias at the onset of kinesin stepping.

One important limitation of the present study is the use of a partial kinesin construct, limited to residues 1–370 of the human kinesin heavy chain, which encompass the motor head, neck linker, and neck helix. However, the full-length kinesin molecule is substantially longer and includes additional structural elements, such as the coiled-coil stalk and cargo-binding domains. These omitted regions may influence kinesin’s conformational dynamics and stepping behavior through long-range mechanical coupling, steric constraints, or altered flexibility. Although the precise impact of these excluded segments remains unclear, their absence could potentially affect the quantitative details of the initial stepping motion observed in our simulations. However, our primary findings are based on the local conformational dynamics of the kinesin motor heads and the neck region within their immediate microtubule-bound context, and are thus expected to remain valid even in the full-length kinesin context. The close agreement between our simulations and previous experimental observations provides strong support for the robustness and reliability of our conclusions.

In this regard, the negatively charged, flexible C-terminal tails (E-hooks) of *α*- and *β*-tubulins, which were also omitted in our simulations, may play an important role. Previous studies have demonstrated that E-hooks enhance kinesin processivity by increasing run length through reduced detachment rates.^49,50^ Although a direct role for E-hooks in the stepping mechanism itself appears limited, their negatively charged nature strongly suggests potential electrostatic tethering interactions with positively charged regions of kinesin, particularly the neck coiled-coil. Indeed, the net positive charge of the neck coiled-coil region (+4 per neck helix, resulting in +8 for the neck coiled-coil) supports the feasibility of such electrostatic interactions, a concept previously illustrated in Figure 4 of Ref. 11. Importantly, these electrostatic tethering interactions are not limited to truncated kinesin constructs; rather, they are likely to persist or even become more prominent in the context of full-length kinesin interacting with an E-hook-containing microtubule lattice. Thus, explicitly incorporating the E-hooks into structural models would be expected to reinforce and further stabilize the neck-region–microtubule interactions we have identified, significantly strengthening our central conclusion that these interactions decisively bias kinesin’s initial stepping motion. Future studies employing more complete kinesin constructs and explicitly modeling the E-hooks will be valuable for refining and extending these insights. Altogether, we believe the current study provides a robust and novel structural framework for understanding the molecular basis of kinesin’s stepping directionality.

In summary, we have presented the first all-atom molecular dynamics study of a dimeric human kinesin-1 on a multi-protofilament microtubule. Our gREST simulations revealed a robust and previously uncharacterized conformation of the neck region that interacts with the microtubule surface. Based on this newly identified conformation, we performed simulations of the initial stepping process. The observed counterclockwise movement of the rear kinesin, consistent with previous experimental findings, highlights the functional importance of neckregion–microtubule interactions in regulating initial stepping directionality. By integrating enhanced sampling techniques, standard molecular dynamics simulations, and a structurally realistic kinesin–microtubule model, our work provides new mechanistic insight into how local structural elements guide directional motion at the onset of kinesin stepping.

## METHODS

### System construction

The simulation system is based on a dimeric human kinesin-1 bound to a microtubule. The kinesin construct includes residues 1–370 of the human kinesin-1 heavy chain, which encompasses the motor head, neck linker, and neck helix. As in many previous studies on human kinesin-1, a cysteine-light variant (C7S, C65A, C168A, C174S, C294A, and C330S)^6^ was used in the present study. The ATP-bound rear head and nucleotide-free front head were modeled based on the X-ray crystal structures reported as PDB entries 4HNA^23^ and 4LNU,^24^ respectively. Missing residues in these structures were modeled using MODELLER. ^22^ Each kinesin motor head was positioned on the microtubule surface by aligning the *αβ*-tubulin heterodimer subunits from the respective PDB entries to the corresponding segments in the high-resolution cryo-EM structure of a three-protofilament microtubule (PDB entry 3J6F^25^). The neck region connecting the two motor domains, for which no high-resolution structural data are available, was modeled as described in the main text.

### Molecular dynamics simulations

All molecular dynamics (MD) simulations in this study were performed using the GENESIS software package^13,14^ on the supercomputer Fugaku. Two different force fields were employed to assess force-field dependence: AMBER ff99SB-ILDN^19,20^ and CHARMM36m^21^ for kinesin and microtubule, and the TIP3P model^51^ was used for water molecules. The kinesin–microtubule complex was placed in a periodic cubic water box, ensuring a minimum distance of 10 Å from the protein surface to the box boundaries. The system was neutralized with K^+^ and Cl*^−^* counterions, and additional ions were added to reach a physiological salt concentration of 100 mM. The full simulation system contained approximately three million atoms. This number was reduced to around two million in the gREST and initial stepping simulations, where certain *αβ*-tubulin subunits were omitted. Before initiating production simulations, the solvated systems were equilibrated in the NPT ensemble (310 K and 1 atm) for 1 ns. A time step of 3.5 fs was employed using a hydrogen mass repartitioning scheme optimized for long time step integration.^52^ Temperature and pressure were regulated using the Bussi thermostat and barostat.^53^ Subsequent production simulations—both gREST and conventional MD—were carried out in the NVT ensemble.

### gREST

For enhanced sampling of the neck region, we employed the generalized replica exchange with solute tempering (gREST) method.^12^ This technique enables selective temperature scaling not only of a specified solute region, but also of specific components of the interaction potential. In the present study, the solute region was defined as the dihedral and nonbonded interaction terms associated with the neck-linker region. Twelve replicas were simulated with effective solute temperatures set to 310, 332, 357, 385, 415, 449, 486, 530, 577, 630, 690, and 760 K, while the remainder of the system—including solvent and non-solute regions—was maintained at 310 K. Replica exchange between adjacent temperatures was attempted every 3000 steps, with the temperature distribution chosen to achieve an exchange acceptance ratio of approximately 0.25. Each replica was simulated for 1 *µ*s under both the AMBER and CHARMM force fields, resulting in a total simulation time of 24 *µ*s. All subsequent analyses were performed on the trajectory sampled at 310 K.

## Data availability

The data that support the findings of this study are available from the corresponding author upon reasonable request. Source data are provided with this paper.

## Acknowledgement

This work was supported by MEXT JSPS Kakenhi (grant numbers 22H02042 (to S.-H.C.), 23H04434 and 24K01996 (to R.I.)). The computer resources were provided by the HPCI system research project (Project ID: hp220225, hp230271, and hp240260).

## Author contributions

S.-H.C. and R.I. designed and initiated the research. S.-H.C. modeled simulation systems, performed simulations, and analyzed data. S.-H.C. and R.I. wrote the manuscript.

## Additional information

**Supplementary information** The online version contains supplementary material available at XXXX.

**Correspondence** and requests for materials should be addressed to S.-H.C. or R.I.

