## Supplementary Information for "Neck-region–microtubule interactions direct counterclockwise stepping of kinesin-1"

### Supplementary Figures

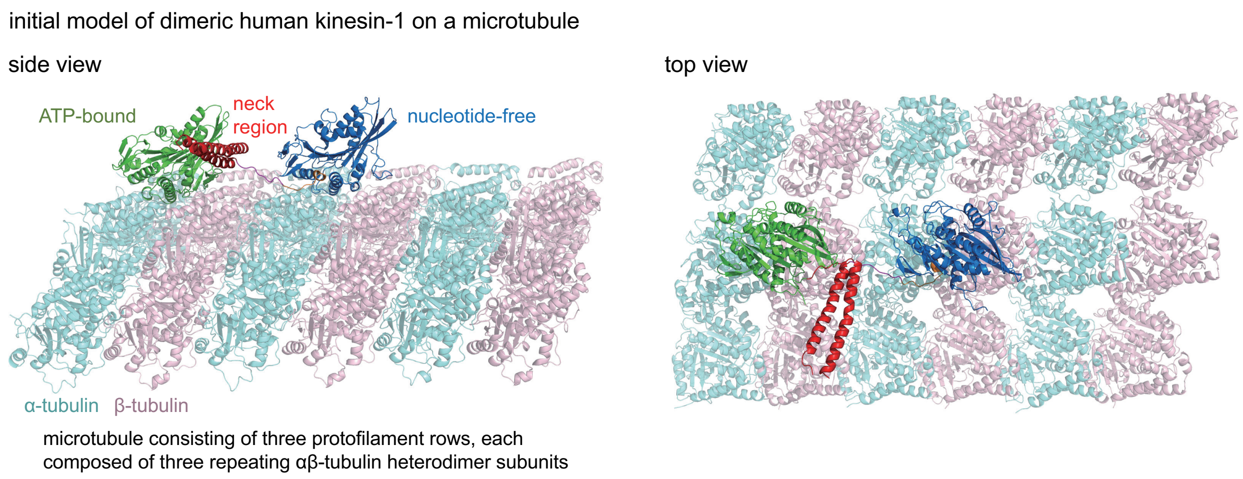


Figure S1: Initial model of dimeric human kinesin-1 bound to a microtubule, shown in side view (left) and top view (right). The modeled neck region is highlighted in red.

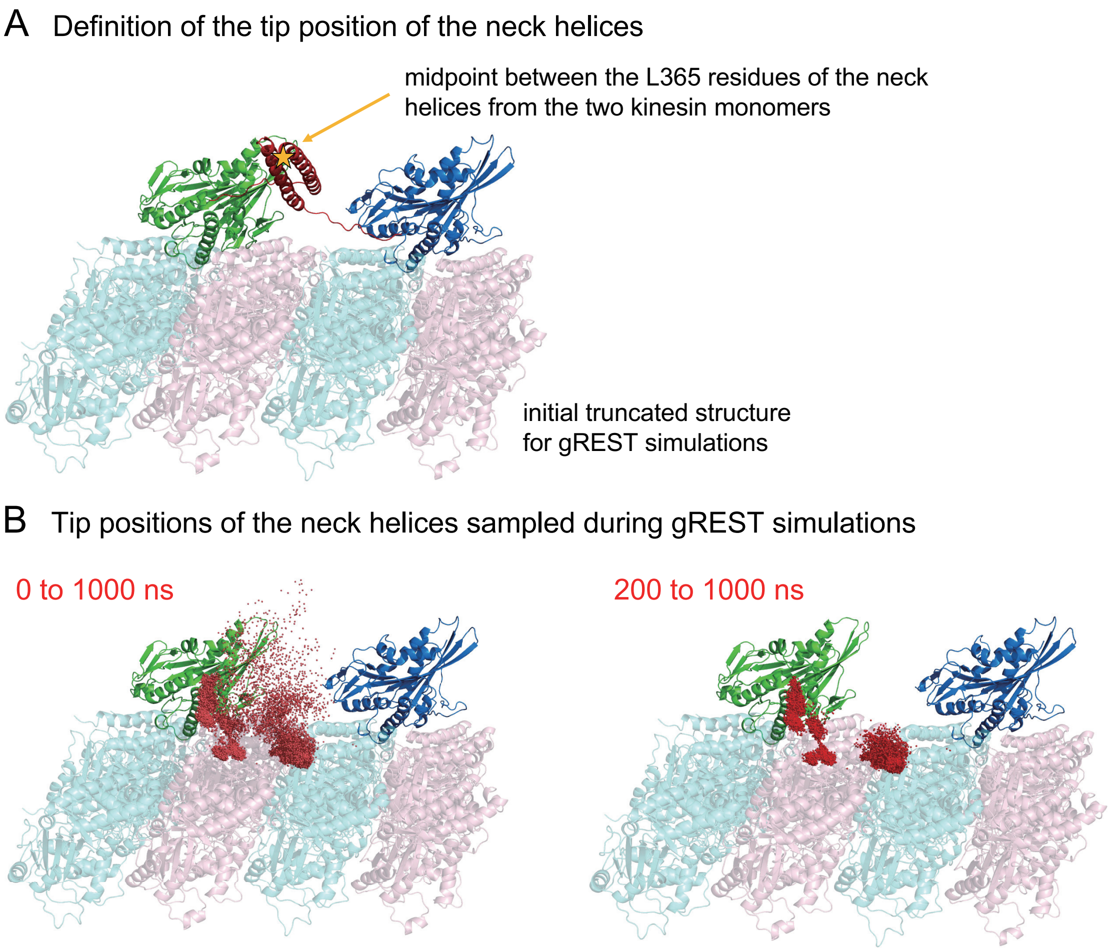


Figure S2: Tip positions of the neck helices sampled during gREST simulations. (A) The tip position for each kinesin dimer conformation is defined as the midpoint between the L365 residues of the neck helices from the two kinesin monomers. The orange star indicates this position for the initial truncated structure used in the gREST simulations. (B) Tip positions sampled over the full course of the gREST simulations (0–1000 ns) and from the later stage (200–1000 ns), excluding the initial 200 ns, are shown as red dots in the left and right panels, respectively.

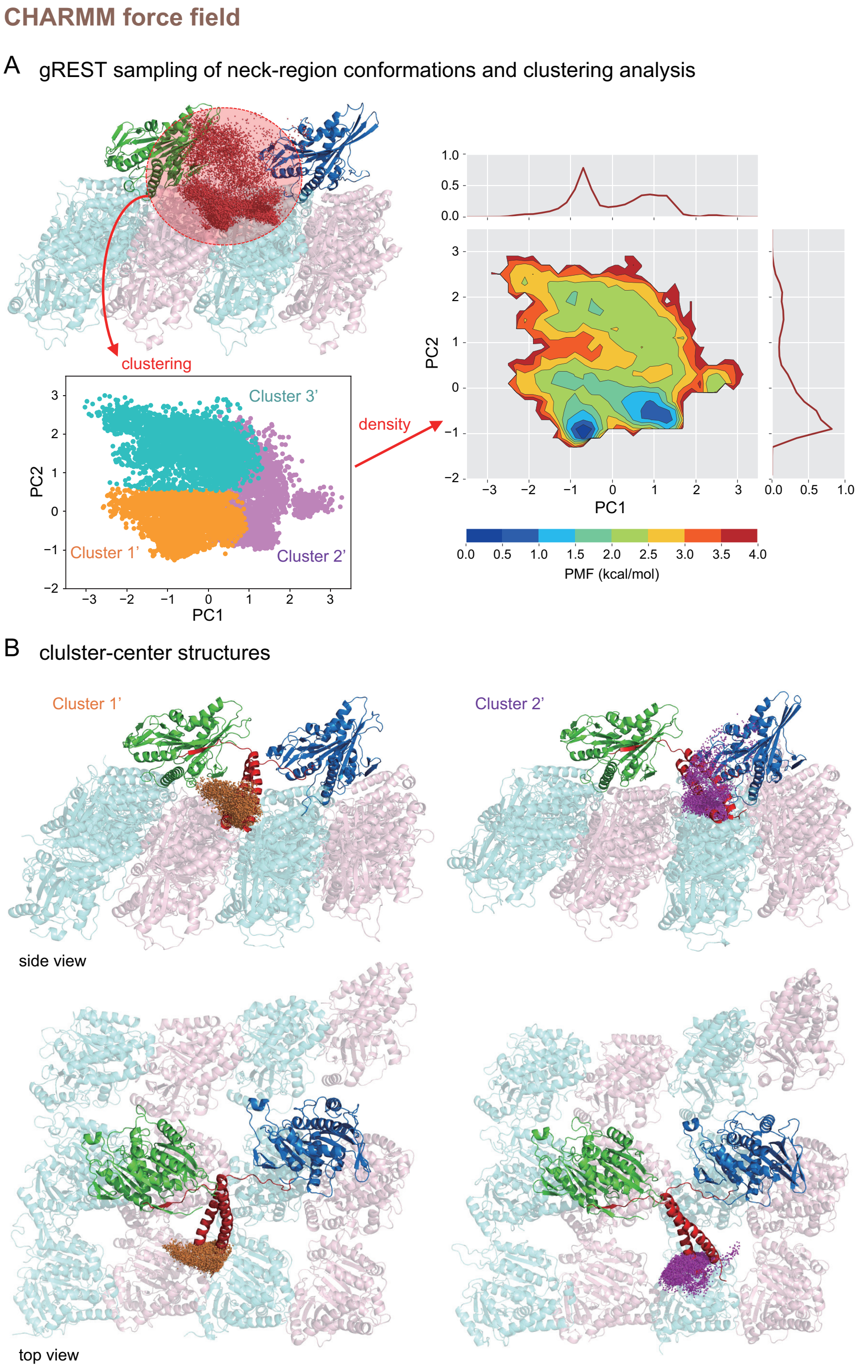


Figure S3: Clustering analysis of neck-region conformations sampled by gREST simulations using the CHARMM36m force field. The same analysis protocol as in Figure 3 of the main text (which used the AMBER ff99SB-ILDN force field) was applied. Two major clusters were identified and labeled as Cluster 1' and Cluster 2' to distinguish them from those presented in the main text. Although an additional cluster (Cluster 3') can be inferred from the clustering results on the principal component (PC) axes, it is not discussed further due to its low population density. The representative structure of the most populated Cluster 1' closely resembles that of Cluster 1 in Figure 3B, highlighting the consistency of the neck-region conformation across different force fields.

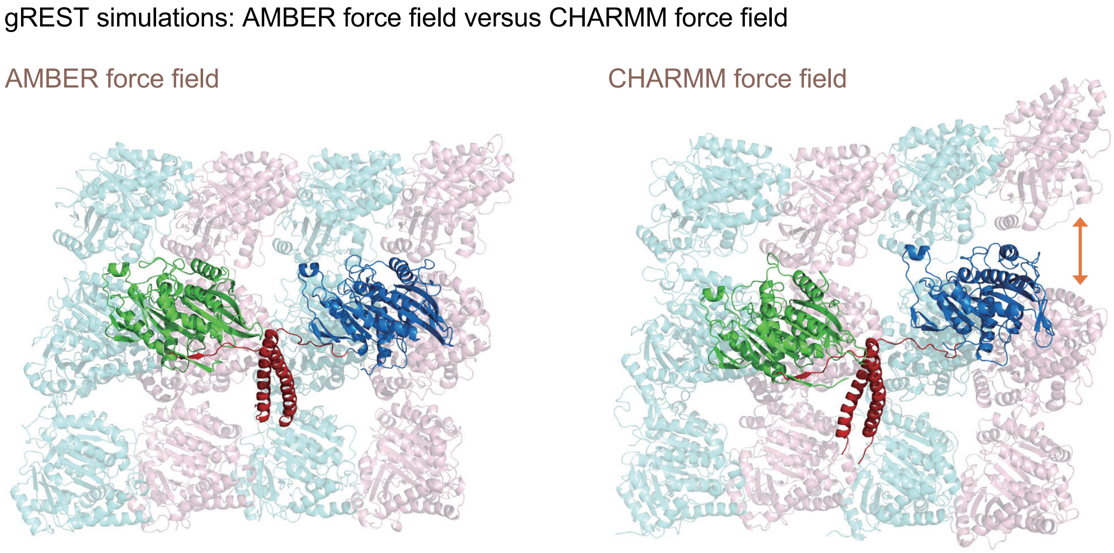


Figure S4: gREST simulations: AMBER force field versus CHARMM force field. Comparison of microtubule structural stability during gREST simulations performed with the AMBER ff99SB-ILDN (left) and CHARMM36m (right) force fields. Each panel shows a representative snapshot from the respective simulation. The AMBER simulation preserved the structural integrity of the multi-protofilament microtubule throughout. By contrast, in the CHARMM simulation, the distance between the central protofilament and one of its neighbors increased markedly, leading to partial disruption of the microtubule lattice.

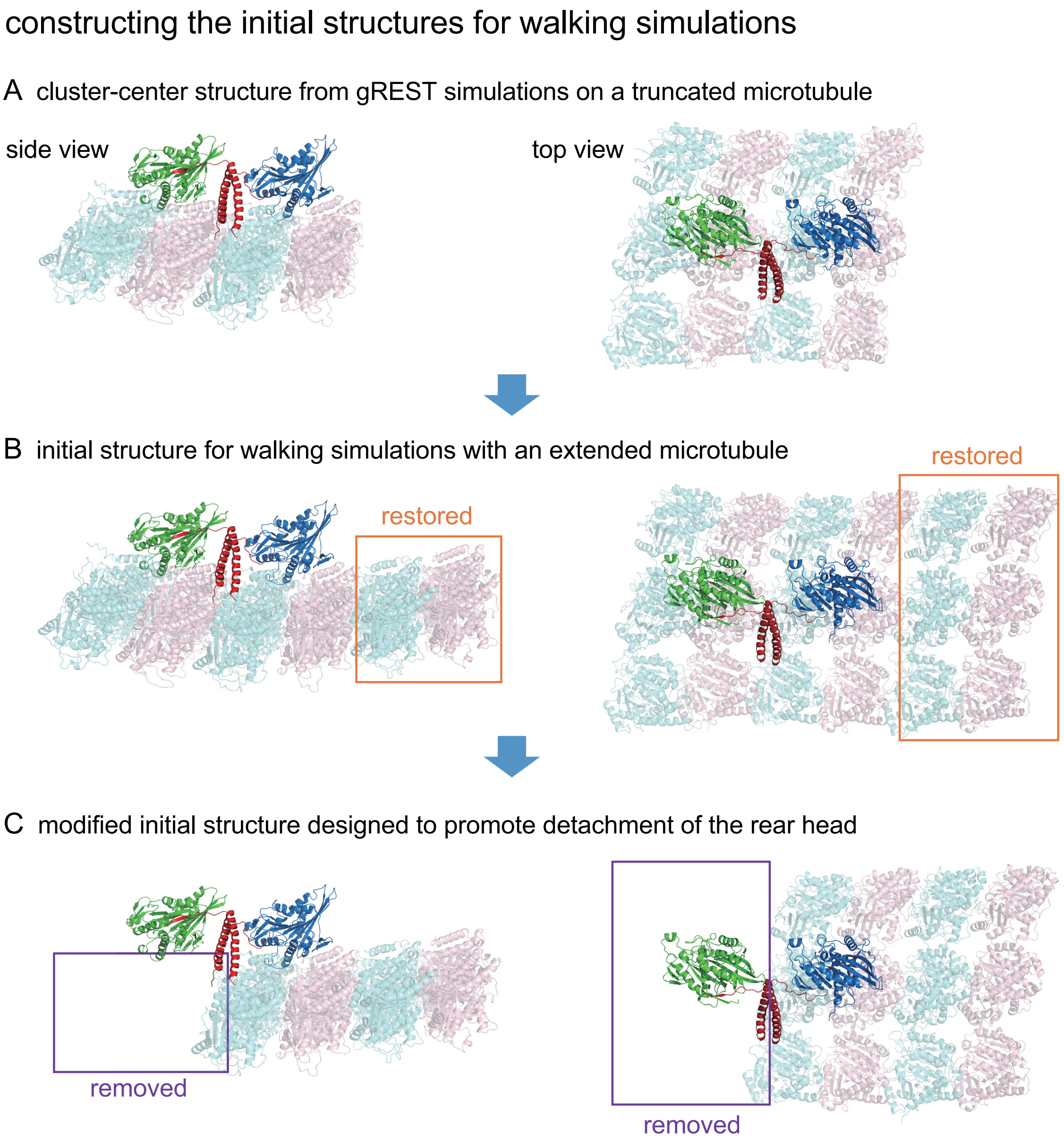


Figure S5: Constructing the initial structures for walking simulations on the microtubule. (A) Cluster 1 structure obtained from the gREST simulations, shown in side and top views, employing a truncated microtubule model. (B) Initial structure for walking simulations with an extended microtubule, in which the distal $\alpha\beta$-tubulin heterodimer subunits removed during the gREST simulations are restored to provide a landing site for stepping. (C) Modified initial structure designed to promote detachment of the rear head, in which the $\alpha\beta$-tubulin heterodimer directly beneath the rear head, along with adjacent subunits from neighboring protofilaments, has been removed.

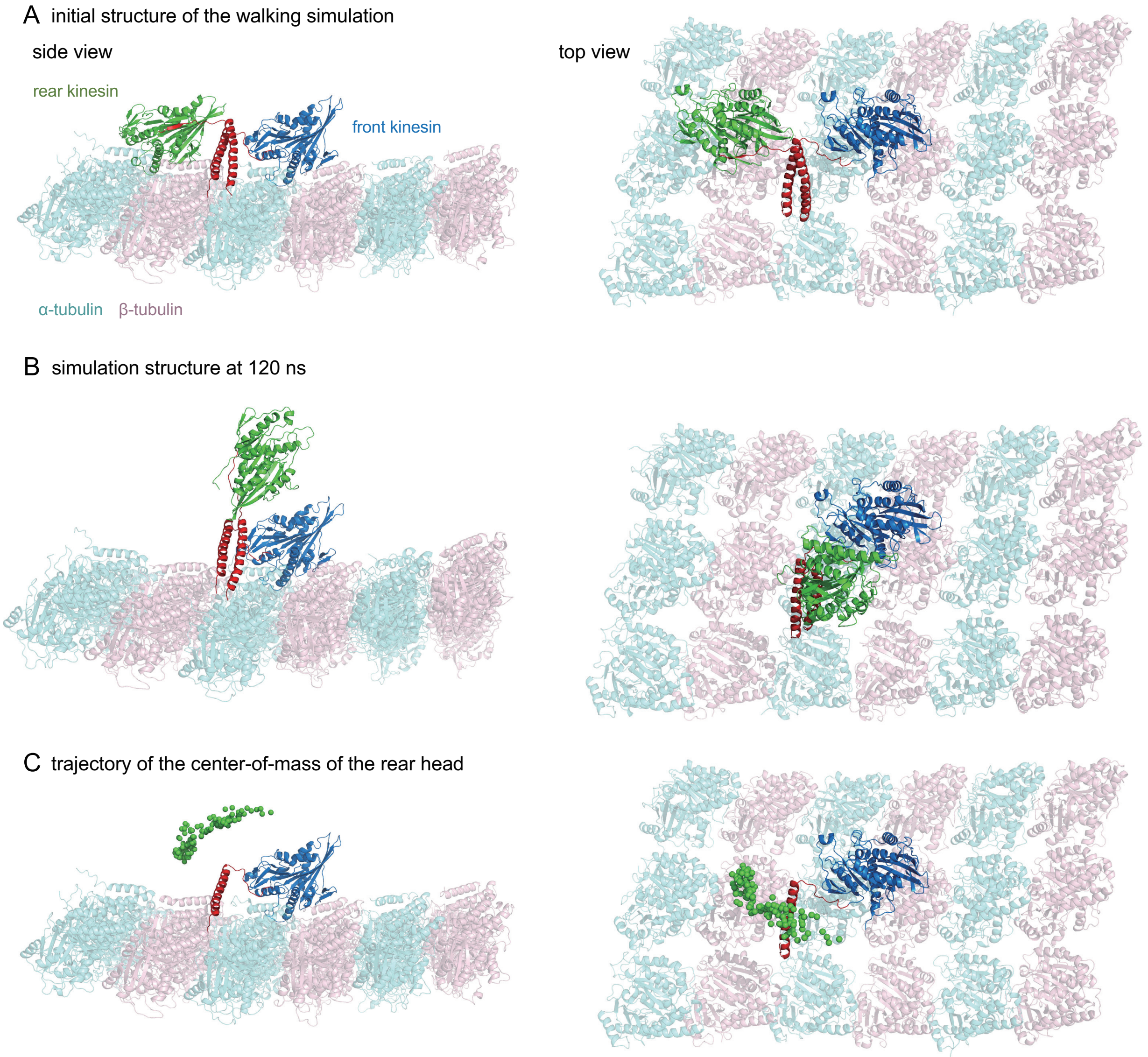


Figure S6: Another representative trajectory of the initial stepping motion of the kinesin dimer on the microtubule. (A, B) Structural snapshots at 0 ns and 120 ns, respectively, from a different simulation starting from the same Cluster 1 conformation. The removed $\alpha\beta$-tubulin subunits have been reintroduced for visualization purposes. (C) Center-of-mass positions of the rear head, sampled every 1 ns from 1 to 120 ns, are shown as green spheres to depict the stepping trajectory.
